# Concurrent contextual and time-distant mnemonic information co-exist as feedback in human visual cortex

**DOI:** 10.1101/2021.06.16.448735

**Authors:** Javier Ortiz-Tudela, Johanna Bergmann, Matthew Bennett, Isabelle Ehrlich, Lars Muckli, Yee Lee Shing

## Abstract

Efficient processing of visual environment necessitates the integration of incoming sensory evidence with concurrent contextual inputs and mnemonic content from our past experiences. To delineate how this integration takes place in the brain, we studied modulations of feedback neural patterns in non-stimulated areas of the early visual cortex in humans (i.e., V1 and V2). Using functional magnetic resonance imaging and multivariate pattern analysis, we show that both, concurrent contextual and time-distant mnemonic information, coexist in V1/V2 as feedback signals. The extent to which mnemonic information is reinstated in V1/V2 depends on whether the information is retrieved episodically or semantically. These results demonstrate that our stream of visual experience contains not just information from the visual surrounding, but also memory-based predictions internally generated in the brain.

**One-Sentence Summary:** Feedback activity in human early visual cortex contains concurrent contextual and time-distant mnemonic information.

## Main Text

Successful navigation through our world requires the efficient integration of sensory inputs with existing knowledge. The predictive processing framework (*1–3*) explains how this integration could take place within the visual domain in the context of a hierarchically organized system. Within this framework, pre-existing knowledge exerts its influence in the form of top-down (i.e., feedback) signals sent from higher to lower levels in the hierarchy; these signals are then contrasted against incoming (i.e., feedforward) information initiated at the retinas. An inherent challenge when studying these dynamics is how to isolate feedback from feedforward signals, since both are assumed to be continuously integrated in all levels (*1*). However, the retinotopic organization of the visual cortex can be utilized for such isolation by physically occluding a portion of the visual field while recording activity from the brain regions that respond to the occluded area. Any modulation of activity in those brain regions, which do not receive any meaningful environmental stimulation, can be taken as a measure of feedback (*4*). Previous research using visual occlusions has shown that mental models of surrounding information, comparable to line drawings, are processed in human V1 and V2 (*5*). However, whether such internal models contain only an extension of low-level visual features of neighboring locations (i.e., extensions of lines) or knowledge about expected scene content (i.e., objects typically found in a scene) has not been investigated so far.

One of the assumptions that is often implicit in research on predictive processing is that feedback signals stem from higher-order representations (e.g., prior knowledge) and that these representations provide information useful to lower-level computations. Here, we investigate whether feedback signals may be constituted by two types of information. Namely, concurrent and mnemonic information (Fig. 1). Concurrent refers to information reaching the visual system as a whole at a given point in time (e.g., the contextual surrounding), but which does not reach specific visual regions directly through feedforward; instead, this information may be relayed via lateral or feedback connections within the visual cortex. In contrast, mnemonic refers to time-distant information that is no longer available from the environment but may be retrieved internally (from stored memory traces, for example). Indeed, if context acts as an effective retrieval cue which accesses a stored memory trace, information contained in that trace can be sent down to lower levels and help to disambiguate the percept based on previous knowledge (*2, 6*).

**Fig. 1.**
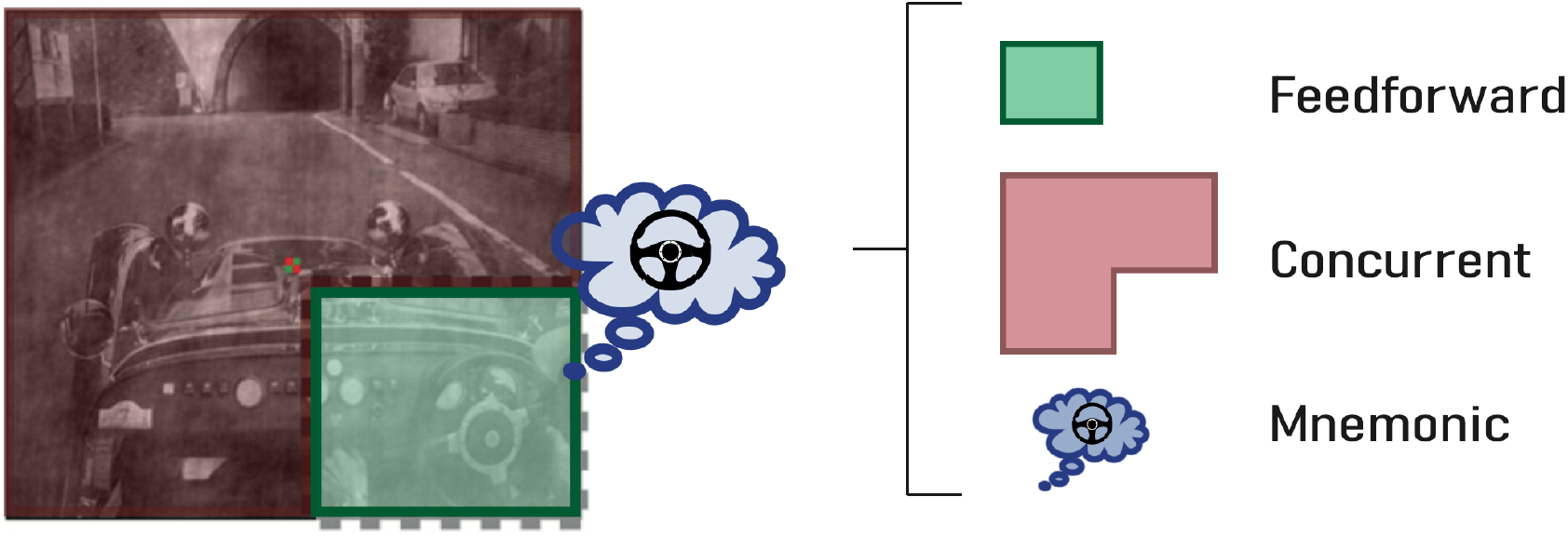
Schematic decomposition of the modulation of brain activity in EVC areas. Green: activity of brain voxels that code for a given portion of the visual field (the steering wheel region) is predominantly modulated by direct sensory information from the eyes (i.e., feedforward information). Red: content that is concurrently presented to other brain regions (i.e., concurrent information) biases activity in voxels in green area via lateral and by providing contextual information. Blue: time-distant information that is not currently presented but that can be accessed through memory (i.e., mnemonic information). These three sources of modulation jointly contribute to perceptual disambiguation.

An important feature of memory traces is the nature of the information that they contain and the neural substrates on which they rely (*7, 8*). Episodic memories, defined as information of past experiences that are bound to a specific spatio-temporal context, are assumed to be rich in perceptual details from the event that generated the episode (e.g., your breakfast this morning) and to rely critically on parietal and medial temporal structures (*9, 10*). Conversely, semantic memories, reflect generalized real-world knowledge acquired through a life-long set of experiences and convey more abstract information (e.g., the general concept of breakfast). Their representations involve a much more distributed network of multimodal regions including posterior (e.g., inferior parietal lobe, middle temporal gyrus and fusiform cortex), medial (e.g., parahippocampal gyri and posterior cingulate gyrus) and prefrontal (e.g., dorsomedial and ventromedial prefrontal cortex) regions (*11*). Both types of memories can serve as sources informing predictions for efficient visual processing. However, the mechanisms underlying memory influences on prediction generation remain unclear.

We used partially occluded stimuli embedded in a memory task to study the content and origin of feedback signals in the early visual cortex (EVC). More specifically, we used functional magnetic resonance imaging (fMRI) to record brain activity while a sample of twenty-nine human participants viewed images depicting a set of scenes. The images were overlaid with a white rectangular patch that hid a target object (Fig. 2A). Participants were instructed to mentally represent the entire scene including the missing object behind the occluder. These missing objects could be accessed either episodically, from a pre-trained set of arbitrary room-object pairs studied 24h before, or semantically, according to participants’ world knowledge of which studied object would fit best to the room (without pre-training; see Fig. 2B for a schematic of the paradigm and Fig. S1 for participants performance). Using functional retinotopy we identify brain voxels that represented the portion of the visual field where we placed the white patch and thus, we isolated activity patterns from non-stimulated regions of the visual cortex. We used multivariate analyses to decode the information content from the activity patterns of these non-stimulated brain voxels. Finally, to assess the involvement of non-visual regions in our task, we performed ROI-based and whole brain connectivity analyses. ROI-based analysis focused on hippocampus (HC) and ventromedial prefrontal cortex (vmPFC) due to their putative role in episodic and semantic retrievals, respectively (*12*); whole brain analysis probed psychophysiological interactions (PPI) to assess changes in functional connectivity between visual regions and the rest of the brain as a function of task condition. Prior to data collection, a pre-registration was created and is available at https://osf.io/va6fc.

**Fig. 2.**
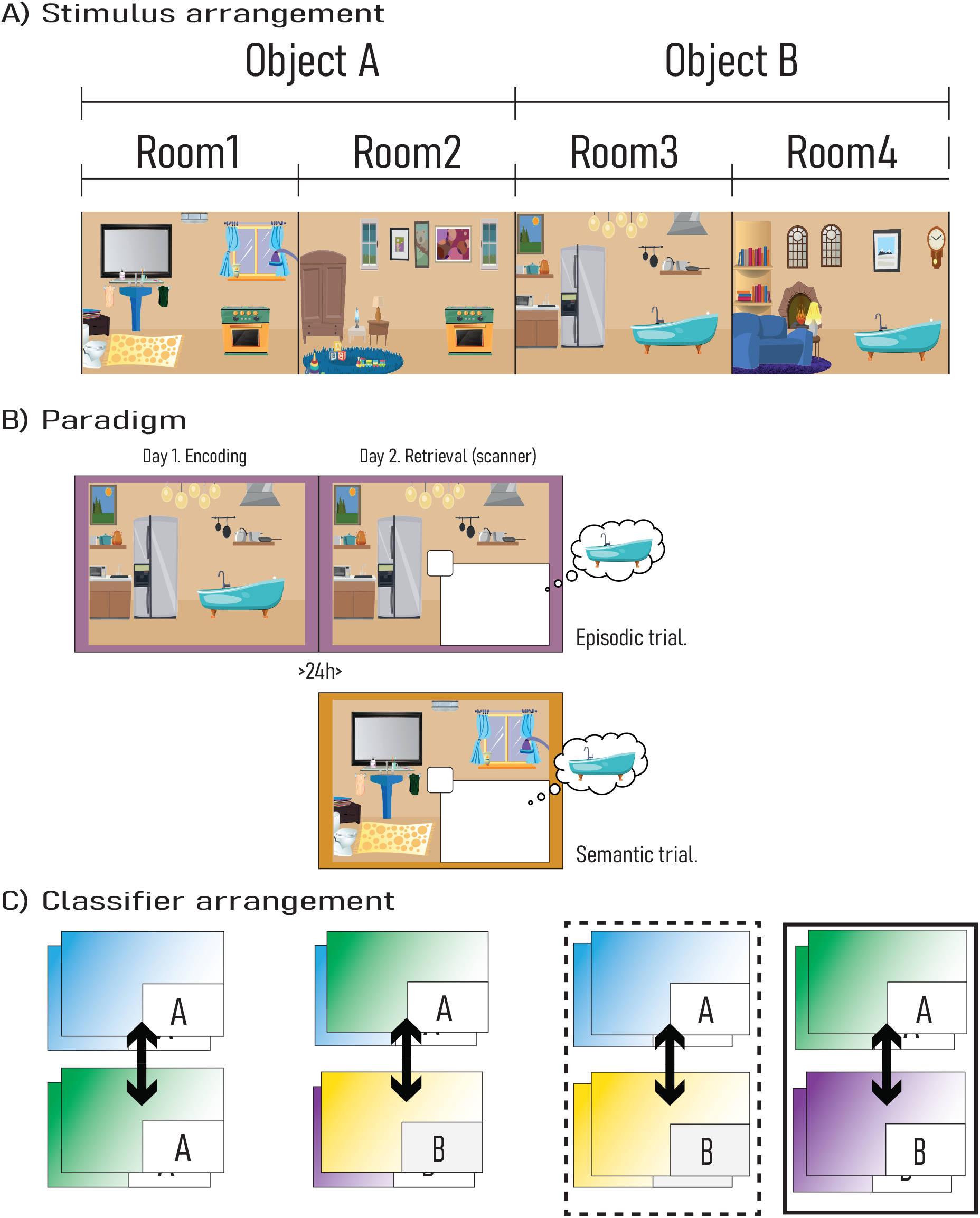
Basic experimental schematics. (**A**) Four example trials of how objects were assigned to rooms. Each object was assigned to two different rooms. (**B**) Experimental paradigm. On episodic trials participants retrieved the object studied on day 1; on semantic trials participants retrieved the object that would semantically fit the room (**C**) Classification set ups. Left-most panel: To test for concurrent information, classifiers were trained and tested to distinguish between different-room trials that shared the to-be-remembered object. Middle-panel: To test for mnemonic content, classifiers were trained and tested to distinguish between different-object trials; each class comprised trials different-room pictures that shared the to-be-remembered object. Right-most panel: To test for the generalization of the reinstated representation, classifiers were trained to distinguish between different-object trials (dashed line); these classifiers were then tested on a different set of pictures that were associated with the same objects as the training set.

### Feedback signals in EVC contain both concurrent and mnemonic information

To test for the presence of feedback signals in non-stimulated brain regions we trained a set of linear support-vector machine (SVM) classifiers on the activity patterns of our regions of interest (ROI) in the EVC. We defined regions of interest using retinotopic mapping to delineate V1 and V2. We confirmed mapped coordinates of occluded regions within V1 and V2 by contrasting mapping conditions (occluded region> surround). Within our ROIs each voxel’s time course captures BOLD activity at that location. We transferred voxel time courses into beta estimates with a Least Squares Separate General Linear Model approach (*13*) to obtain trial-specific estimates. We used the voxel beta estimates to train SVMs to classify condition information and cross-validated classifier performance by training on three runs and testing on the fourth run. The first set of SVMs classified between different rooms paired with the same objects (Fig. 2C). Since memory content was kept constant across trials, classification performance in occluded V1 and V2 ROIs would index the presence of concurrent room information. Our classifiers performed above chance level (50%) for both episodic and semantic trials in V1 and V2 (episodic trials: z=4.69 and z=4.69; semantic trials: z=4.58 and z=4.62, all ps<.001, one-sided Wilcoxon tests; full summary of results can be seen in Table S1). Hence, although the EVC regions from which we decoded did not receive any feedforward signals, information about the concurrent surround (i.e., the different rooms) was still being represented (Fig. 3). This pattern of results replicates and extends previous findings using visual occlusions by showing that concurrent visual information is a constituent part of the feedback activity in V1/V2.

**Fig. 3.**
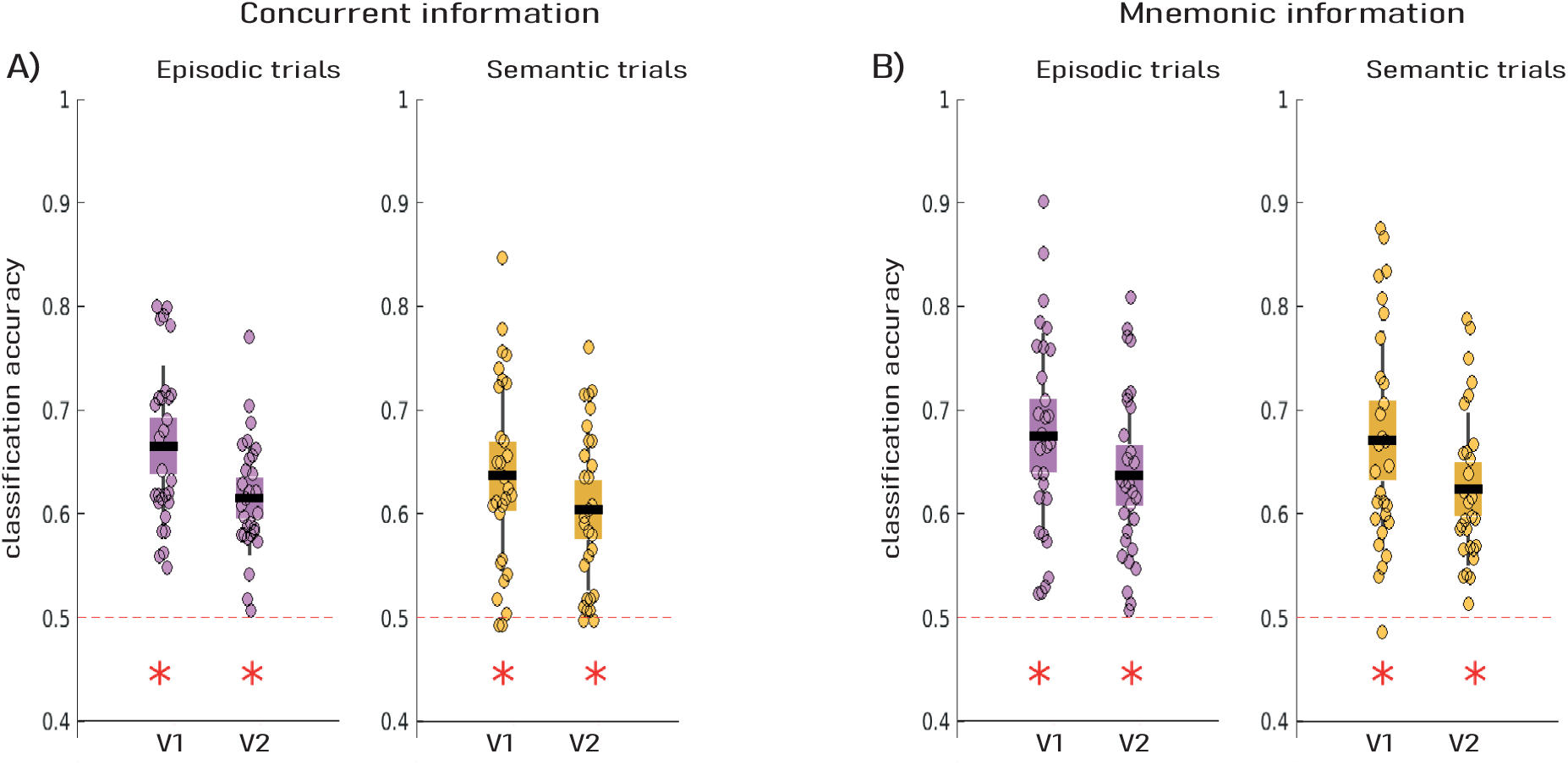
Classifier performance in functionally delineated EVC ROIs. Cross-validated classification accuracy is shown for concurrent (**A**) and mnemonic (**B**) information classifiers. Each dot represents one participant, and the colour of the box plots denotes the retrieval type (i.e., episodic or semantic). Red asterisks indicate above chance (50%) performance.

In addition to the modulation of EVC activity by concurrent feedback signals, we assessed the effect of memory-driven feedback by classifying between different object retrievals (Fig. 2B). We collapsed trials in which the retrieved object was the same despite having different room information. The results revealed the presence of memory information in the occluded portions of V1/V2 (episodic trials: z=4.69 and z=4.70; semantic trials: z=4.67 and z=4.69, all ps<.001 in one-sided Wilcoxon tests). These results complement those above by showing that object memory information can be observed in non-stimulated regions of the EVC (see *Supplementary Materials* for spatial specificity tests and cross-classification).

Our SVM classification results suggest that concurrent and mnemonic information can be decoded from non-stimulated regions in V1/V2. To estimate the contribution of these types of information to the feedback signal, we used Representational (Dis)Similarity Analysis (*14*). We computed separated Representational Dissimilarity Matrices (RDMs) for each ROI in the EVC and created two model RDMs - one for each type of information (Fig. 4A). The model RDM for concurrent information had maximal values in pairs of trials that shared the same room, whereas the model RDM for mnemonic information had maximal values for pairs of trials that shared the same object. Neural and model RDMs were correlated using Spearman’s rho (Fig. 4B and *Supplementary Text*). Since we were not interested in the shared variance across models, we isolated the unique contribution of each model to the group level RDM using variance partitioning (*15, 16*). With this approach we could determine the unique fraction of variance in the ROI RDM that is explained by the concurrent or the mnemonic model RDM. We conducted three linear regressions (one for each type of information separately and another one for both together) with model RDMs as predictors and the ROI RDMs as predictands. Then, we inferred the amount of unique variance explained by each model by subtracting the explained variance of each single regression from the explained variance of the combined regression (Fig. 4C). As expected, the concurrent model uniquely explained a significant portion of the variance in both ROIs across both trial conditions. This confirms the modulation of activity in occluded regions by information that is concurrently fed back from surrounding non-occluded regions. On the other hand, once the variance explained by the concurrent model was accounted for, the mnemonic model was still able to explain a significant portion of the remaining variance in EVC for episodic trials (Fig. 4D; all ps<.001). For semantic trials, however, the memory model was unable to capture any variance that was not already explained by the concurrent model (Fig. 4D; all ps>.1). By removing the contribution of the concurrent information, this result indicates that memory-specific reinstatement was observed in occluded regions of EVC but that feedback predictions contain mnemonic information only when it is retrieved episodically, not semantically.

**Fig. 4.**
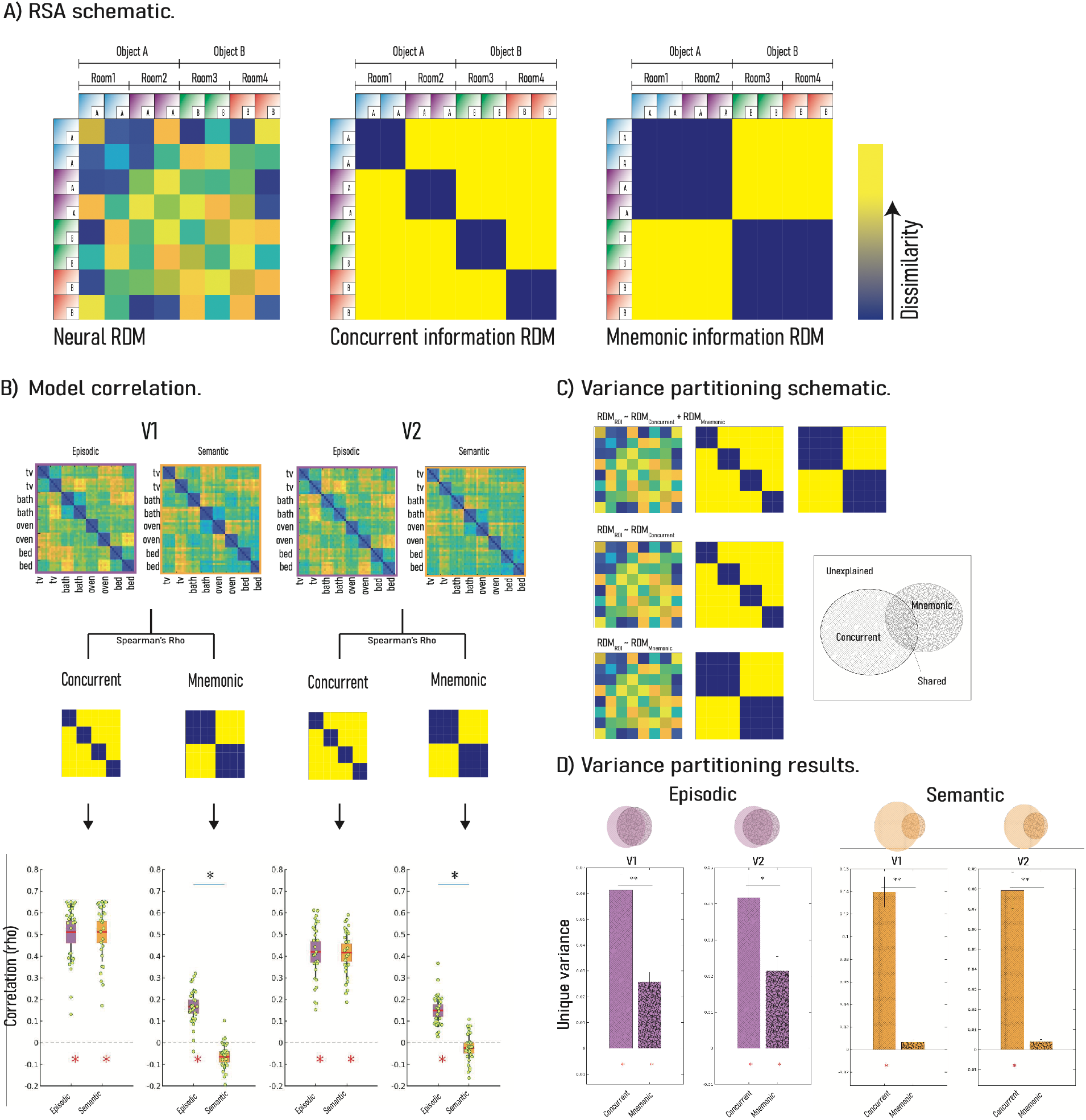
Representational Similarity Analysis in occluded regions of the EVC. (**A**) Schematic of how ROI and model RDMs were constructed (see also *Main Text*). For neural RDMs, correlational distances (Pearson) were computed for every pair of trials; for the concurrent RDM, maximal values (yellow) were used same-room trials and minimal values (blue) for different-room trials; for the mnemonic RDM, maximal values (yellow) were used for same-object and minimal values (blue) for different-object trials. (B) Correlation of each model RDM was with the ROI RDMs for episodic and semantic trials. (C) Schematic representation of the variance partitioning analysis. Three regression models were fitted with the ROI RDM as the predictand and the different models as predictors. The first regression (top-panel) included both models as predictors, the second and third regressions included only one of the models (middle and bottom panels). By comparing the variance explained in each one of the regressions, we can infer the fraction of the uniquely explained by each predictor. The Venn diagram depicts the rationale for the variance partitioning analysis for the concurrent and the mnemonic models. (**C and D**) Variance partitioning results for episodic and semantic trials. The colour and the pattern of the bars denote the retrieval type and the model tested, respectively. Red asterisks indicate non-zero uniquely explained variance (p<.05); black asterisks indicate significant differences in the amount of explained variance by each model (* = p<.05; ** = p<.001). Note that, when the variance explained by the concurrent model was removed, the mnemonic model was only able to explain a significant portion of variance for episodic but not for semantic trials.

To validate the sensitivity of our paradigm to capture representational similarities among semantic trials and since our target memories were real-world objects, we computed a new RDM in the object-selective cortex (LOC; see Supplementary Materials) and assessed an object-specific reinstatement index for each participant. The index was computed by subtracting the dissimilarity measures of different scene-same object trials from those of different scene-different object trials (Fig. 5A). Since this index is computed directly from distance measures in the RDMs, it has a straightforward interpretation in terms of representational change: any above zero value indicates that the retrieval of the same object in two different rooms increased the representational similarity in that ROI. We observed significant object reinstatement in LOC for both episodic and semantic trials, both p<.001, suggesting that objects retrieved semantically as well as episodically were represented in LOC. Moreover, when contrasted with a two-sided Wilcoxon test, the index did not differ between trial conditions, z=0.638, p=.523, suggesting that object reinstatement in LOC was equivalent in both episodic and semantic conditions. To enable a direct comparison between LOC and the EVC, we extended this index to the occluded portions of V1/V2. As can be seen in Fig. 5B, and in line with the variance partitioning results, we observed object reinstatement in occluded V1/V2 for the episodic condition (V1: z=4.65, V2: z=4.69, ps<.001). In contrast, there was no reinstatement in either ROI for semantic trials (V1: z=−3.84, V2: z=−1.70, ps>.05). These results were further confirmed by a significant interaction in a repeated-measures ANOVA with ROI and trial type as within-participants factors, F(3,196)=12.448, p<.001. This comparison confirms that our participants were able to access and represent the objects retrieved in both conditions, but only episodically retrieved information reached the EVC as a prediction signal.

**Fig. 5.**
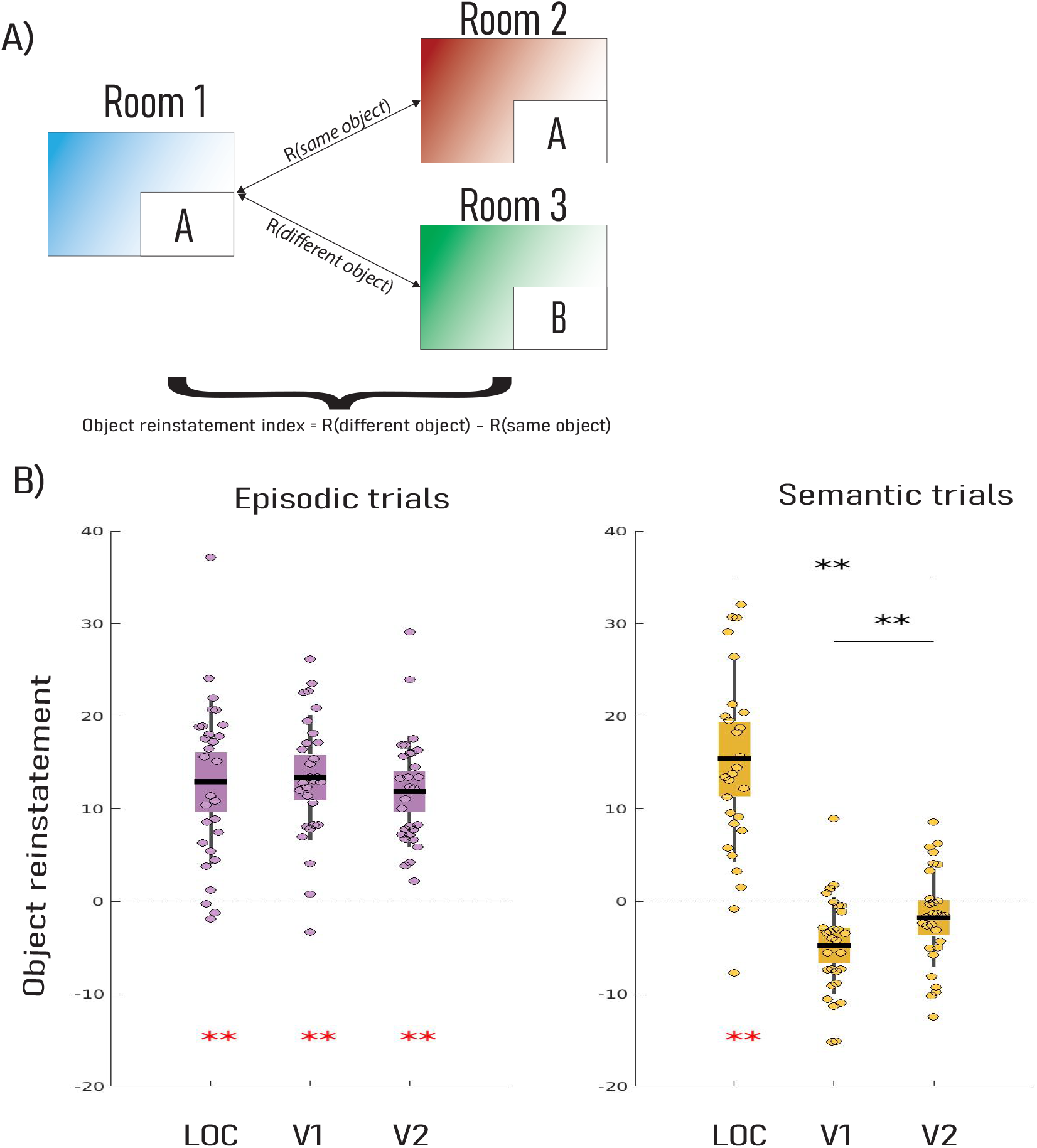
Object-specific reinstatement index. **(A)** An object-specific reinstatement index was computed by assessing the average dissimilarity between same-object pairs (top arrow) and subtracting it from the average dissimilarity between different-object pairs (bottom row). Non-zero values in this index represent increase in similarity during the retrieval of the same object. **B**) Object reinstatement in visual cortex ROIs. Red asterisks indicate significant object reinstatement (<.05); black asterisks indicate significant differences in the amount of reinstatement in each ROI (* = p<.05; ** = p<.001). While object reinstatement was similar in LOC for both episodic and semantic retrievals, in V1/V2 we saw object reinstatement only in episodic but not semantic trials.

### Different potential sources for predictions during semantic and episodic trials

Higher-order mnemonic representations are assumed to exist across distributed neural regions, depending on whether they are episodic or semantic in nature and are potential sources for time-distant content carried by in feedback signals. To locate potential source regions used trial-level classifier decision values as an index of how distinct the neural pattern was for a given object class, with patterns laying further from the decision boundary being more representative exemplars of their class (*17*). These decision values from V1/V2 occluded regions were correlated with activity in HC and vmPFC, due to their respective putative roles in episodic and semantic memories (*12*). Subject-wise correlations were Fisher Z-transformed and tested against zero with a one-sided Wilcoxon test. Consistent with previous research highlighting the involvement of HC in episodic memory tasks and as anticipated in our pre-registration, we observed that the distinctiveness of the representations in visual cortex was correlated with HC activity in episodic (R = 0.0297, z=2.30, p=.021) but not in semantic trials (R = −0.0296, z=−1.09, p<.05; Fig. 6A). This correlation supports the involvement of HC in reinstating representational patterns in the visual cortex (28). In sharp contrast, while we had hypothesized a positive correlation for vmPFC during semantic trials, we did not find such correlation for either memory condition (both p>.05). Although the involvement of vmPFC in semantic retrievals has been extensively reported before (*11, 12*), the lack of object reinstatement in visual cortex in semantic trials might have prevented any correlation with activity in distant regions (see *Supplementary Text* for full brain exploratory analyses).

**Fig. 6.**
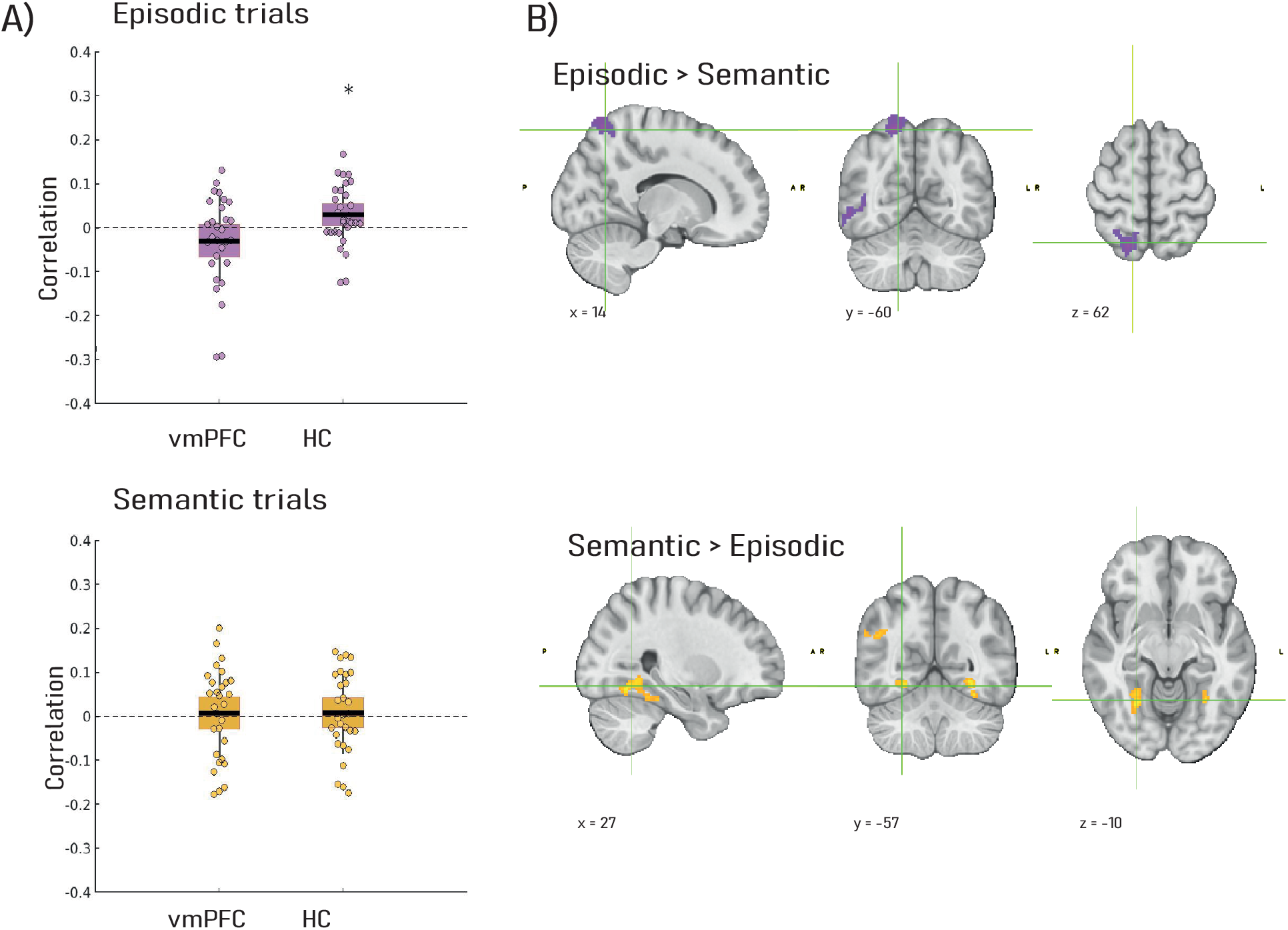
Relating activity in distant brain regions to feedback signals. (**A**) Correlation between trial-level classifier decision values in occluded regions and beta estimates in non-visual ROIs. (**B**) Whole-brain PPI analysis with LOC as seed region and retrieval type as psychological variable. The results revealed increased functional connectivity between LOC and right parietal during episodic trials and between LOC and bilateral fusiform for semantic trials.

Finally, the whole-brain PPI analysis included the two memory conditions as main psychological modulators of functional connectivity. Since object reinstatement was equivalent in LOC for both memory conditions, we selected LOC as our seed region to be able to establish a fair comparison between conditions (see *Supplementary Text* for parallel analysis with V1). The results of this analysis revealed increased functional connectivity between LOC and a subregion of the right posterior parietal cortex during episodic trials (p<.05, cluster corrected). In contrast, semantic trials increased functional coupling between LOC and fusiform gyrus (p<.05, cluster corrected). These results reflect that object reinstatement engages a different set of regions based on whether the reinstatement arises via an episodic or a semantic route (Fig. 6B; see *Supplementary Table* S2 for a summary of all clusters with increased connectivity). Although PPI does not allow causal inferences, the differential pattern of connectivity for our two memory conditions point towards brain regions that are potentially crucial for memory-based prediction generation.

## Discussion

Visual processing largely relies on the integration of prior knowledge with sensory inputs. Within the predictive processing framework, it is proposed that incoming feedforward information is integrated with feedback predictions across all levels of the processing hierarchy (*1, 2*). For such integration to take place, feedback signals need to carry informational content. Our study demonstrated that one of the main sources for this content is mnemonic information by showing that 1) feedback predictions can be detected in V1 and V2 in the human EVC (*4*), 2) these feedback predictions can carry both concurrent information about the immediate surrounding context and mnemonic information that is formed in the past, 3) the extent to which mnemonic information is fed down to EVC critically depends on the retrieval route through which that information is accessed and 4) episodic and semantic retrievals differentially engage dorsal and ventral brain regions, respectively. Taken together, our results advance our understanding on the role that memory systems play in informing visual predictions. While in the prediction literature, priors are often based on acquired relationships between basic low-level features (e.g., line orientations), we show that higher-order mnemonic information can inform priors as well. These results bring together predictive processing and memory systems, two major fields in cognitive neuroscience that are in principle closely related but are often considered in isolation.

### Predictions carry different informational content

Occlusions happen constantly in our daily life and object recognition from partially occluded inputs is one of the challenges that our perceptual system needs to overcome. In addition, perceptual disambiguation is not restricted to partial occlusions, since any perceptual experience can be understood essentially as a form of probabilistic inference about the most likely stimuli to have caused the perceptual input (*6, 18*). In order to efficiently and accurately disambiguate sensory inputs, our brains need to integrate contextual information with pre-existing knowledge. Here, we show that both types of information can be detected in feedback signals from the earliest of visual cortex regions (i.e., V1 and V2). The co-existence of concurrent contextual content and time-distant mnemonic information enables us to disambiguate perceptual input and to avoid inappropriate inferences driven by memory alone. A rather ambiguous input (e.g., seeing a drill in a poorly lit room) can be clarified based on its surrounding information (e.g., it is placed on a workshop table next to other tools (*6*)) and/or from memories of previous experiences (e.g., last time you did some repairs in your basement). In such conditions, and in agreement with our present results, the clarifying information will be constituted by a combination of concurrent surrounding information and of episodic memory content. Although the example above describes a situation in which both concurrent and mnemonic information point to the same concept (e.g., drilling tool), it is possible to think of situations where they might point to different concepts (e.g., a hairdryer to dry some wet painting). Whether in agreement or disagreement, the presence of both types of signals in the same brain structure is a pre-requisite for integration and for appropriate inferences about the most likely object being perceived (*6, 18*).

Finally, it is also worth noting that, while our results show that a concurrent vs. mnemonic parcellation can be performed on feedback signals, concurrent information is very likely to be in turn a mixture of low-level perceptual features (e.g., lighting conditions, predominant colours, etc.) and mid-level categorical information (e.g., objects commonly found in a basement). Although this distinction cannot be addressed by our current results, we argue that they lay the ground for future studies specifically targeting the content of concurrent feedback signals in EVC.

### Feedback predictions from semantic and episodic differ in content and neural substrates

Following Tulving’s seminal definition, episodic memories are thought to be richer in perceptual details than semanticized memories which, by means of abstraction, have lost representational distinctiveness in favor of generalizability (8, 21). However, the neural instantiation of that difference had not yet been fully explored in the context of feedback signals (*17, 19*). Interestingly, while activity in LOC has been linked to global object feedforward processing (*20*), V1 and V2 are particularly fine-tuned to low level features in incoming stimuli. It is conceivable that this differential preference for varying levels of features in feedforward signals might extend to reinstated features from mnemonic representations in the feedback pathway. Accordingly, we show that the reinstatement of an object’s features, under episodic and semantic trials, was similar in higher visual regions, namely LOC, but differed greatly in V1 and V2. Namely, during episodic trials, the object-specific pattern was successfully reinstated in non-stimulated subregions of V1 and V2; in contrast, there was no reinstatement during semantic trials. The difference in reinstatement between higher and lower visual regions and, more specifically, the lack of reinstatement during semantic trials in V1 and V2 is very likely to reflect a difference in the preference for low-level perceptual features. Note that although our SVM classifiers were successful to distinguish between different object retrievals also in semantic trials, the variance partitioning results suggest that classifier accuracy was most likely entirely driven by concurrent information.

Finally, our connectivity results indicate that semantic and episodic reinstatements also rely on different brain networks. The posterior parietal cortex is considered to be part of a wider hippocampal memory network, which is particularly engaged when accessing temporally bound memories (*21*). On the other hand, while semantic representations are assumed to be widely distributed over the cortex, activity in some regions such as the fusiform gyrus have been shown to be involved in semantic tasks involving object selection (*22*) and categorization (*11*). The requirements of our task closely mimic these processes. In episodic trials, participants reinstated the unique room-object combinations learnt in the study session to retrieve the appropriate object; in semantic trials participants had to correctly identify the room category shown (e.g., bathroom) and to select the semantically fitting object (e.g., a bathtub) among four possible candidates. Accordingly, our results revealed that functional connectivity between LOC and the posterior parietal cortex was enhanced during the retrieval of scene-object pairs. In contrast, we observed that LOC changed its functional coupling from dorsal to ventral regions (i.e., fusiform gyrus) during semantic retrievals. Due to the lack of directionality in PPI analysis we cannot ascertain whether LOC was modulated by distant activity or vice versa. However, the differential crosstalk between an area engaged in active object reinstatement and a set of memory related networks is compatible with the idea that top-down predictions stem from different higher order (memory) representations.

The results encapsulated in this paper uncover an important piece of the puzzle of the role that memory plays in generating predictions. The retrieval of pre-existing information can undeniably serve as a source mechanism to inform prediction but, is then prediction just memory retrieval? It is most likely not. As discusses above, prediction involves information from different sources, and it is through the integration of these pieces that top-down modulations can be exerted (*1, 2*). In other words, predictive processing, from the generation of the predictions themselves to their continuous updating, involves several processes one of which is memory retrieval. Does this mean that memory retrieval serves the only purpose of prediction generation? This might be at least partially true. Here we focused on declarative memory, namely episodic and semantic memories, where the subjective experience of retrieval can be detached from action and therefore it can serve an ulterior purpose (e.g., mental imagery, cf. (*23*)). Other types of memory such as non-declarative procedural memories can be even more strongly related to prediction, especially in the motor control domain. The automatized retrieval of well-learnt actions and skills is so closely intermingled with predictive dynamics (*24*) that the two might not be separable. It would be more accurate, therefore, to say that (declarative) memory and prediction are two different but interactive systems that jointly contribute to render an efficient and seemingly continuous perceptual experience.

## Acknowledgments

We thank members of the LISCO lab (PI: YLS), Carlos Gonzalez-García and Catarina Sanches-Ferreira for their helpful comments on this research.

## Funding

Provide complete funding information, including grant numbers, complete funding agency names, and recipient’s initials. Each funding source should be listed in a separate paragraph.

Examples:

European Research Council Starting grant ERC-2018-StG-PIVOTAL-758898 (YLS).

Jacobs Foundation Research Fellowship JRF 2018–2020 (YLS)

German Research Foundation Project ID 327654276, SFB 1315, “Mechanisms and Disturbances in Memory Consolidation: From Synapses to Systems” (YLS)

Goethe Research Academy for Early Career Researchers - Fokus A/B program (JO)

## Author contributions

Conceptualization: JO, YLS

Data curation: JO, IE

Formal analysis: JO, IE

Funding acquisition: YLS

Investigation: JO, IE

Methodology: JO, YLS, JB, MB, LM

Project administration: JO, YLS

Software: JO, IE

Supervision: YLS, LM

Visualization: JO, IE

Writing – original draft: JO

Writing – review & editing: JO, YLS, JB, MB, IE, LM

## Data and materials availability

- **Stimulation and analysis scripts:** https://github.com/ortiztud/feedbes
- **Raw data:** https://openneuro.org/git/1/ds003691
- **Unthresholded Z-maps from PPI:** https://neurovault.org/collections/UAFOYKRZ/

