## supplementary text for "Concurrent contextual and time-distant mnemonic information co-exist as feedback in human visual cortex"

**This PDF file includes:**

Materials and Methods
Supplementary Text
Fig. S1
Tables S1 to S2

### Materials and Methods

Registration

Prior to data collection, a registration was created at https://osf.io/va6fc. Any deviation from the registration is indicated in the corresponding section.

Participants

Thirty healthy young adults (18 female; age: mean = 24.41, sd = 2.87) were recruited through advertisements placed across the three campi of the Goethe University in Frankfurt. They all gave written informed consent to participate in this study, in accordance with the local ethics committee. In exchange for participation, they received either course credits (for Psychology majors) or honorarium (for all other majors). All participants had normal or corrected-to-normal vision. One of the participants had to be excluded due to excessive movement during three out of the four functional runs in the scanning session.

Stimuli and materials.

Previous studies using partially occluded images have used real world photographs as stimuli (*1–3*). When using this kind of stimuli, the location of the occluder bears no relation to the structure of the image content itself, thus resulting in seemingly random cuts in the picture. These random cuts generate a set of line-stops right at the edge of the occluder (e.g., tree trunks or traffic lights) that randomly vary from photograph to photograph but that are constant for all the repetitions of the same photograph. As a consequence, differences between photographs could be measured based on mere interpolations from line stops without the need for any top-down influence. Since our aim was to study mnemonic content, we created a novel stimulus set that did not include any meaningful line stops. This set consisted of sixteen cartoon images depicting common indoor locations and four cartoon images of commonly used objects. Importantly, all pictures were adjusted so that the location that the occluder will cover included only one line (i.e., a straight line that separated the floor from the wall) that was held constant across all pictures. All materials are available at https://github.com/ortiztud/feedbes.

Our stimulus set was divided into two subsets. For the episodic subset, eight scene-object pairs were built so that the two elements had a minimal semantic relation (e.g., "TV" in a "bathroom"). The remaining eight scenes were used for the semantic subset, and they were paired with objects that had a strong semantic relationship (e.g., "oven" in a "kitchen"). Critically, every object was paired with two scenes in each retrieval condition (Fig. 2A). Every participant was assigned to a given combination of objects and scenes and these combinations were counterbalanced across participants so that across the entire sample every object was seen in every possible scene.

During the tasks outside the scanner, stimuli were displayed using a 60 Hz monitor (resolution: 1680 x 1050, full HD) placed approximately 60cm away from participants' heads. During the tasks inside the scanner, stimuli were displayed using a 60 Hz monitor (resolution 1920 x 1080, full HD) placed 162cm away from participants' eye (eye to mirror + mirror to monitor distance). Stimuli spanned 16.4º x 12.1º of visual angle.

Procedure.

*Day 1:*

Learning phase: Participants were presented with five learning blocks during which the pairs from the episodic set were shown sequentially in a computer screen and repeated ten times. Participants were instructed to memorize as many details as possible from the entire scene and to pay special attention to the target object (i.e., the object that was placed on the bottom-right corner of the scene). At the end of each block, knowledge about the scene-object pairings was tested with two tests. Namely, memory for the object identity was measured with a four-alternative forced choice test with the scene as cue and the four objects as options; memory for the precise object position was measured by presenting the object misplaced and asking participants to place it back into its original position using the arrow keys in the keyboard. After completing the learning blocks, participants were given a printed version of the scenes with a white patch occluding the target area and they were asked to draw the target objects from memory. The between-blocks memory tests and the drawing task were included here to ensure that participants formed distinctive and detail-rich memories for each pair.

After the learning phase, participants performed one block of the object retrieval task (equivalent to one scanning run; see below) to familiarize themselves with the timing and the structure of the task. The only difference between this familiarization block and the scanner blocks is that a trial-by-trial vividness rating was performed to further promote the rich visualization of perceptual features of the objects. On every trial, participants were asked to report their subjective vividness of the retrieved object on a four points scale.

Finally, at the end of the session, one last learning block, identical to those at the beginning, was conducted as a refreshment of the studied stimuli.

*Day 2 (24h after Day 1):*
Pre-scan phase: Right before entering the scanner, participants were briefly shown the scenes in the semantic set. They were asked to remember the objects studied on day 1 and to mentally select the one that would best fit each scene (i.e., those that would be typically found in each location in the real world) and to verbally report it. If they selected the wrong object, the correct answer was provided by the experimenter. This only happened once across the entire sample due to a speeded response from one participant.

Scanning phase: In order to avoid fatigue and to prevent unwanted movements, we distributed our scanning sequences over two sessions with a 5 to 10 minutes break in between; participants could use the toilet, stretch and walk to prevent neck and shoulders stiffness. Both scanning sessions included one T1 weighted anatomical image, two object retrieval runs and one visual mapping run. In addition, the first session included a high-resolution scan of the hippocampal area and an extra visual mapping run, and the second session included an extra functional run not used for the current project.

Occlusion task: This task consisted of two types of trials: episodic and semantic. On episodic trials participants were presented with the scenes studied during the learning phase on day 1; on semantic trials, participants were presented with a new set of scenes. In both trial types, a white patch occluded the bottom right corner of the image thus hiding the target object. Upon encountering an episodic trial, participants were asked to vividly retrieve the corresponding studied object from the previous day (episodic retrieval); when presented with a semantic trial, participants were asked to mentally select, among the four possible objects, the one that would fit the scene in the real world (semantic retrievals). Trial order transitions were optimized (*4*) to allow for maximum separation of trial types.

Each of the sixteen scene-object combinations was shown six times along each run and each trial lasted 4 seconds with an inter-trial interval of 2 seconds. During the 4 seconds of the trial, the image flashed at a 5Hz frequency. For the entire duration of the trials a special type of fixation cross was used to minimize eye movements (*5*).

Retinotopic mapping: To map our participants' visual cortex activity to on-screen positions we used standard stimulation procedures of eccentricity and polar angle mapping. For eccentricity mapping a contrast-reversing checkerboard expanding ring was displayed at the centre of the screen; one full cycle of expansion lasted 56 seconds and a total of 9 cycles were shown. For polar angle mapping, a contrast-reversing checkerboard rotating wedge was presented centred in the screen; the wedge rotated clockwise to cover the entire screen after 64 seconds for a total of 8 complete rotations.

Target mapping: We used contrast-reversing flashing checkerboards (flashing frequency 5Hz) to functionally locate the voxels in V1 and V2 responding to our target region (i.e., bottom-right corner of the screen). Since previous studies have shown that successfully representing objects can recruit foveal voxels (*6*), we also included a checkerboard patch that spanned 2º of visual angle from the centre of the screen. To allow for maximum separation of the different conditions, each pattern was repeated 6 times following a 12s on/12s off block design. A central fixation cross was shown during the entire duration of the task. To ensure fixation, participants were asked to monitor the colour of the fixation cross and to press a button every time that it changed colour; colour changes were randomly presented during the on periods a maximum of three times.

Post-scan phase: Participants performed a memory test (identical to the one in the learning phase) that checked for scene-object associations (both for episodic and semantic pairings). In addition, in the case of the episodic trials, we checked for correct positioning of the objects in the scene.

<u>MRI data acquisition</u>

MRI data was acquired using a 3 Tesla MR scanner (SIEMENS Prisma) with a 32-channel head coil at the Brain Imaging Center (BIC) of the Goethe University (Frankfurt, Germany). In both scanning sessions, a 3D anatomical scan (3D MPRAGE; 1 x 1 x 1 mm resolution; iPAT factor: 2) was acquired. In addition, a high-resolution Turbo Spin Echo scan (TSE; 0.4 x 0.4 x 2 mm resolution; TE= 16ms; TR= 6500ms) was acquired during the first session; the 206mm field of view was placed over each participant's hippocampus (HC) by first locating the left HC and aligning the shorter axis of the field of view to the long axis of the HC. Blood oxygen level-dependent (BOLD) signals were measured with an echo-planar imaging sequence (EPI; TE= 38ms; TR= 800ms; resolution= 2 x 2 x 2 mm; iPAT factor= 8; flip angle= 52º field of view=208 mm; 72 axial slices; phase encoding direction= AP). Finally, two extra sets of 5 volumes of the EPI sequence were acquired in each phase encoding direction to allow for susceptibility distortion correction.

<u>MRI data processing</u>

*Preprocessing*
All structural and functional MRI images, apart from the TSE and EPIs from the retinotopic mapping run, were preprocessed using fMRIPREP 20.0.1 (*7*). A boilerplate text released under a CC0 license describing preprocessing details can be found below. For further information about the pipeline, see fMRIPREP´s documentation. Functional scans from the retinotopic mapping run were slice time corrected, 3D motion corrected and temporally filtered (high-pass filtered at 0.01 Hz and linearly detrended) using Brain Voyager 21.4 for Linux (Brain Innovation).

*Start of the boilerplate>>>*
Results included in this manuscript come from preprocessing performed using *fMRIPrep* 20.1.0 (Esteban, Markiewicz, et al. (2018); Esteban, Blair, et al. (2018); RRID:SCR_016216), which is based on *Nipype* 1.4.2 (Gorgolewski et al. (2011); Gorgolewski et al. (2018); RRID:SCR_002502).

*Anatomical data preprocessing*
A total of 2 T1-weighted (T1w) images were found within the input BIDS dataset. All of them were corrected for intensity non-uniformity (INU) with N4BiasFieldCorrection (Tustison et al. 2010), distributed with ANTs 2.2.0 (Avants et al. 2008, RRID:SCR_004757). The T1w-reference was then skull-stripped with a *Nipype* implementation of the antsBrainExtraction.sh workflow (from ANTs), using OASIS30ANTs as target template. Brain tissue segmentation of cerebrospinal fluid (CSF), white-matter (WM) and gray-matter (GM) was performed on the brain-extracted T1w using fast (FSL 5.0.9, RRID:SCR_002823, Zhang, Brady, and Smith 2001). A T1w-reference map was computed after registration of 2 T1w images (after INU-correction) using mri_robust_template (FreeSurfer 6.0.1, Reuter, Rosas, and Fischl 2010). Volume-based spatial normalization to two standard spaces (MNI152NLin2009cAsym, MNI152NLin6Asym) was performed through nonlinear registration with antsRegistration (ANTs 2.2.0), using brain-extracted versions of both T1w reference and the T1w template. The following templates were selected for spatial normalization: *ICBM 152 Nonlinear Asymmetrical template version 2009c* [Fonov et al. (2009), RRID:SCR_008796; TemplateFlow ID: MNI152NLin2009cAsym], *FSL's MNI ICBM 152 non-linear 6th Generation Asymmetric Average Brain Stereotaxic Registration Model* [Evans et al. (2012), RRID:SCR_002823; TemplateFlow ID: MNI152NLin6Asym],

*Functional data preprocessing*
For each of the 6 BOLD runs found per subject (across all tasks and sessions), the following preprocessing was performed. First, a reference volume and its skull-stripped version were generated using a custom methodology of *fMRIPrep*. Head-motion parameters with respect to the BOLD reference (transformation matrices, and six corresponding rotation and translation parameters) are estimated before any spatiotemporal filtering using mcflirt (FSL 5.0.9, Jenkinson et al. 2002). BOLD runs were slice-time corrected using 3dTshift from AFNI 20160207 (Cox and Hyde 1997, RRID:SCR_005927). A B0-nonuniformity map (or *fieldmap*) was estimated based on two (or more) echo-planar imaging (EPI) references with opposing phase-encoding directions, with 3dQwarp Cox and Hyde (1997) (AFNI 20160207). Based on the estimated susceptibility distortion, a corrected EPI (echo-planar imaging) reference was calculated for a more accurate co-registration with the anatomical reference. The BOLD reference was then co-registered to the T1w reference using flirt (FSL 5.0.9, Jenkinson and Smith 2001) with the boundary-based registration (Greve and Fischl 2009) cost-function. Co-registration was configured with nine degrees of freedom to account for distortions remaining in the BOLD reference. The BOLD time-series

(including slice-timing correction when applied) were resampled onto their original, native space by applying a single, composite transform to correct for head-motion and susceptibility distortions. These resampled BOLD time-series were referred to as *preprocessed BOLD in original space*, or just *preprocessed BOLD*. The BOLD time-series were resampled into standard space, generating a *preprocessed BOLD run in MNI152NLin2009cAsym space*. First, a reference volume and its skull-stripped version were generated using a custom methodology of *fMRIPrep*. Automatic removal of motion artifacts using independent component analysis (ICA-AROMA, Pruim et al. 2015) was performed on the *preprocessed BOLD on MNI space* time-series after removal of non-steady state volumes and spatial smoothing with an isotropic, Gaussian kernel of 6mm FWHM (full-width half-maximum). Corresponding "non-aggresively" denoised runs were produced after such smoothing. Additionally, the "aggressive" noise-regressors were collected and placed in the corresponding confounds file. Several confounding time-series were calculated based on the *preprocessed BOLD*: framewise displacement (FD), DVARS and three region-wise global signals. FD was computed using two formulations following Power (absolute sum of relative motions, Power et al. (2014)) and Jenkinson (relative root mean square displacement between affines, Jenkinson et al. (2002)). FD and DVARS are calculated for each functional run, both using their implementations in *Nipype* (following the definitions by Power et al. 2014). The three global signals are extracted within the CSF, the WM, and the whole-brain masks. Additionally, a set of physiological regressors were extracted to allow for component-based noise correction (*CompCor*, Behzadi et al. 2007). Principal components are estimated after high-pass filtering the *preprocessed BOLD* time-series (using a discrete cosine filter with 128s cut-off) for the two *CompCor* variants: temporal (tCompCor) and anatomical (aCompCor). tCompCor components are then calculated from the top 5% variable voxels within a mask covering the subcortical regions. This subcortical mask is obtained by heavily eroding the brain mask, which ensures it does not include cortical GM regions. For aCompCor, components are calculated within the intersection of the aforementioned mask and the union of CSF and WM masks calculated in T1w space, after their projection to the native space of each functional run (using the inverse BOLD-to-T1w transformation). Components are also calculated separately within the WM and CSF masks. For each CompCor decomposition, the *k* components with the largest singular values are retained, such that the retained components' time series are sufficient to explain 50 percent of variance across the nuisance mask (CSF, WM, combined, or temporal). The remaining components are dropped from consideration. The head-motion estimates calculated in the correction step were also placed within the corresponding confounds file. The confound time series derived from head motion estimates and global signals were expanded with the inclusion of temporal derivatives and quadratic terms for each (Satterthwaite et al. 2013). Frames that exceeded a threshold of 0.5 mm FD or 1.5 standardised DVARS were annotated as motion outliers. All resamplings can be performed with *a single interpolation step* by composing all the pertinent transformations (i.e. head-motion transform matrices, susceptibility distortion correction when available, and co-registrations to anatomical and output spaces). Gridded (volumetric) resamplings were performed using antsApplyTransforms (ANTs), configured with Lanczos interpolation to minimize the smoothing effects of other kernels (Lanczos 1964). Non-gridded (surface) resamplings were performed using mri_vol2surf (FreeSurfer).

Many internal operations of *fMRIPrep* use *Nilearn* 0.6.2 (Abraham et al. 2014, RRID:SCR_001362), mostly within the functional processing workflow. For more details of the pipeline, see the section corresponding to workflows in *fMRIPrep*'s documentation.

<<< *End of boilerplate*

*ROI definitions*

Early visual cortex ROIs. Data from each participant's retinotopic mapping run was used to delineate V1 and V2 using linear cross-correlation of eight polar angle steps and eight eccentricity steps. Data from the target mapping run was fit using a general linear model (GLM) with one predictor per condition (i.e., periphery target area, periphery surround, fovea target area, fovea surround) and the resulting beta estimates were used to select ROIs that responded to the peripheral patch (periphery target area > periphery surround) and to the foveal patch (fovea target area > fovea surround). A conjunction map was computed by masking the contrast maps with the V1 and V2 ROIs obtained from functional retinotopy. At the end of the procedure, we obtained four ROIs (i.e., V1 periphery, V2 periphery and V1 fovea and V2 fovea) representing V1 and V2 voxels that responded to the occluded areas in the periphery and the fovea.

Hippocampus ROIs. From each participant's TSE scan a hippocampal mask in native space was extracted using the automatized software ASHS (*8*).

Atlas based ROIs. Neurosynth (*9*) was used to obtain metanalytic activation maps for the desired brain regions. The searches for "vmpfc", and "LOC" output activation maps based on 151 and 226 studies, respectively. These activation maps were thresholded to a z-score of 5, registered into each subject's native space and turned into binary masks using FSL's built in functions.

Note that only the delineated subregions of V1 and V2 were restricted to occlusion and the rest of the masks were used in their entirety.

*Generalized linear model*

For our multivariate analysis, single-trial beta estimates were obtained by modelling BOLD time course with a series of Generalized Linear Models (GLM) using the Least Square Separate method (*10*, *11*) with LSS-16. The GLM for a given trial contained a total of 32 regressors: one for the onset of the trial, 16 regressors for the onsets for the 16 different scenes, 6 regressors for head motion (3 for displacement and 3 for rotation), 3 regressors for global, WM and CSF intensity and 6 regressors for eye movements (3 for displacement and 3 for rotation). A total of 96 GLMs per run were computed for every participant and only the single-trial estimates were used further.

*Multi-voxel Pattern Analysis (MVPA)*

Single-trial beta estimates for each ROI were used to train and test our classifiers using a leave one run out cross-validation scheme. Three different types of binary classifiers were used (see Figure 2). In order to obtain a measure of concurrent information, the classifiers were trained and tested with different scenes paired with the same objects (e.g., scene 1-object A vs scene 2-object A). To get a measure of mnemonic information, we grouped together different scenes paired with the same objects (e.g., scene 1-object A + scene 2-object A) and labelled those as a function of the object that was being retrieved (e.g., object A vs object B). Finally, in order to get a measure of the transference of object information between retrievals from different scenes, we tested our classifiers with different scenes paired with different objects (e.g., scene 1-object A vs scene 2-object B) and tested them on a different subset of scenes that were paired with the same objects as the train set (e.g., scene 3-object A vs scene 4-object B). The results of the binary classifiers were then averaged for each classifier types to get a single estimate of each type for each participant. All classification analyses were performed with The Decoding Toolbox (*12*).

*Representational Similarity Analysis (RSA)*
RSA analyses were not part of the original registration since they were suggested by one of the authors as a complementary approach to the SVM classifiers once data collection had already started.

Model correlation and variance partitioning: Two ideal binary RDMs were simulated to predict what the neural RDMs should look like in the presence of concurrent or memory information. These model RDMs had zeros in the same scene-same object cells and ones in the rest of the matrix, for the concurrent model, and zeros in all same object cells (i.e., including same-scene and different scene) and ones in the rest of the matrix, for the mnemonic model (see Figure 4); in these model RDMs, zero represents minimal dissimilarity and one represents maximal dissimilarity. Since the resulting model RDMs overlap in their predictions about the same scene-same object trials, when computing the second order RSA with the object model, those cells were not included. Finally, in order to assess the unique contribution of each model RDM to explaining the ROI RDMs, a variance partitioning method was applied (*13*). A group-level RDM was computed by averaging individual RDMs and separate regression models were fit using as regressors either each model separately or both models; then, the unique contribution of each model was computed by subtracting the explained variance ($r^2_{Adj}$) for the other model in isolation from the estimate of both models combined. Significance testing on the fractions was performed by running 1000 permutations of the cells in the ROI RDM and comparing the resulting distribution with the original value.

Object specific index: Single-trial beta estimates for each run were used to compute a Representational Dissimilarity Matrix -RDM- (*14*) for each ROI using The Decoding Toolbox (*12*) and all subsequent analysis were performed with homebrew scripts available at https://github.com/ortiztud/feedbes. RDMs capture the pattern of correlations among the single-trial multivoxel patterns in each ROI with higher values indicating less correlation (i.e., more representational distance). To obtain a distance measure for each object, an object-specific reinstatement index was computed by averaging the correlational distance (1 – Kendall Tau) across trials for different scene-same object pairs and subtracting that from the average distance for different scene-different object pairs (see Figure 5). This index reflects the extent to which retrieving the same object from a different picture decreases the representational distance. The object-specific reinstatement index was average across objects to obtain a single value per ROI per participant.

*Psycho-Physiological Interaction (PPI)*
All PPI analyses were conducted with FEAT (FMRI Expert Analysis Tool) Version 6.00, part of FSL (FMRIB's Software Library, www.fmrib.ox.ac.uk/fsl) (*15*). For 29 participants we performed four PPI analyses, one for each seed region and each contrast direction of episodic and semantic trials, i.e., $\text{PPI-V1}_{\text{EPISEM}}$, $\text{PPI-V1}_{\text{SEMEPI}}$, $\text{PPI-LOC}_{\text{EPISEM}}$, and $\text{PPI-LOC}_{\text{SEMEPI}}$.

In addition to the nuisance regressors that had been described above for the multivariate GLM, our PPI models included three additional regressors. The first regressor (PHYS) represents the physiological signal of the corresponding seed region, the second regressor (PSYCH) codes for the respective contrast of conditions, and the third regressor (PHYS*PSYCH) is the interaction between physiological signal and the psychological condition.

PHYS: For every subject and every run, the average timeseries of both seed ROIs was extracted with FSL's function fslmeants. The resulting vector was entered into the corresponding first-level regression model as a regressor that represents the physiological variable.

PSY: In order to partial out any changes in connectivity that might be driven by main effects of the task, we included in all PPI analyses two psychological regression vectors (PSY_A-B and PSY_A+B) for mean-centering purposes. For PPI-$V1_{EPISEM}$ and PPI-$LOC_{EPISEM}$, PYS_A-B coded episodic trials as 1 and semantic trials as -1, whereas for PPI-$V1_{SEMEPI}$ and PPI-$LOC_{SEMEPI}$ the coding was reversed. PSY_A+B coded both, episodic and semantic trials, as 1. All psychological regressors were convolved with a standard gamma hemodynamic response function (HRF) before they were entered into the PPI models.

PHYS*PSY: The interaction regressor of interest was created by multiplying the demeaned timecourse of the seed region, i.e., PHYS, with the mean-centered psychological vector A-B, i.e., PSY.

All data were high-pass filtered with a cut-off of 100 s. For the first level analysis, three contrasts were specified, i.e. [1 0 0], [0 1 0], and [0 0 1]. All contrasts were computed voxelwise for the four runs that each subject had completed during the experiment. The resulting estimates were passed on to the second level mixed effect analysis (FLAME, FMRIB's Local Analysis of Mixed Effects), to combine the results across runs. Lastly, second level estimates were brought up to the third level between-subjects group analysis, resulting in the final functional connectivity maps. Group level maps were cluster corrected to an alpha value <.05.

*Parametric modulation by decision values*
The decision values from the object classifier were used to parametrically modulate our task regressors in a mass-univariate full brain analysis (*16*). The trial-by-trial decision values were convolved with an HRF and de-meaned before introducing them into the GLM. Two beta maps were obtained for the parametric modulation of the episodic and semantic retrievals. The contrast analysis for episodic > semantic and for semantic > episodic did not reveal any cluster of voxels that survived a cluster correction of p<.05.

### Supplementary Text

Behavioural analysis

Participants' responses from the memory tests at the end of each learning block are shown in Figure S1. Responses from three participants were not recorded, so the total sample for behavioural data is twenty-six. As expected, performance for both the scene-object pairs and for the object position improved over blocks and was maintained in the last block after the scanner training task. The average vividness judgement of each participant during the scanner training task is also shown in Figure S1.

Average responses to the post-scan memory tests are also shown in Figure S1 (bottom panel).

Spatial specificity: Periphery vs. fovea

In line with previous literature showing that foveal voxels can be recruited to accurately represent objects shown in the periphery of the visual field (23), we were also able to measure mnemonic information for our objects from the foveal subregions of V1 and V2 (episodic trials: $z=4.69$, $p<.001$ and $z=4.69$, $p<.001$; semantic trials: $z=4.58$, $p<.001$ and $z=4.69$, $p<.001$, one-sided Wilcoxon signed rank tests). The spatial specificity of the mnemonical content was assessed by contrasting performance on the peripheral ROIs with the foveal ROIs. Separately for episodic and semantical trials, classification accuracy scores for mnemonic information of each participant were tested with a repeated-measures analysis of variance (ANOVA) with ROI (v1 periphery, v2 periphery, v1 fovea and v2 fovea) as within-participants factors. As expected from the original positioning of the objects within the images, the results revealed a main effect of ROI, $F(3,84)=17.958$, $p<.001$ and $F(3,84)=25.17$, $p<.001$, with classifiers performing significantly worse when trained on foveal voxels and peripheral voxels in V1: $F(1,28)=4.39$, $p<.001$ and $F(1,28)=4.98$, $p<.001$, for episodic and semantic sets, respectively and V2: $F(1,28)=4.95$, $p<.001$ and $F(1,28)=7.05$, $p<.001$, for episodic and semantic, respectively. Classification was significantly different from chance in all four ROIs (all $p<.001$); the full summary of results can be seen in Table S1.

Cross-classification.

To get a measure of transfer between retrievals from different cues, we set up a cross-classification scheme in which we trained our classifiers in a subset of different room-different object trials and tested them in a different subset of trials that maintained the same object association structure. In contrast with previous classification analysis, cross-classification did not differ from chance in either memory condition (all $p>.05$), suggesting that the retrieved patterns learned by our classifiers were specific to given room-object combinations.

Model RDM correlation.

Neural and model RDMs were correlated using Spearman's rho. Fischer Z-transformed individual correlation scores were computed and compared against zero with one-sided Wilcoxon tests. The results confirmed the presence of strong concurrent information in all ROIs for both trial types (all $ps<.001$). More interestingly, the mnemonic model was correlated only with ROI RDMs of episodic trials (Fig. 4B) thus suggesting that pattern reinstatement in EVC only took place when object memories were accessed through an episodic route, but not through a semantic route.

Psychophysiological interaction.

Contrasting episodic with semantic trials, we found a cluster of voxels located at the right superior LOC that was functionally stronger connected to both seed regions during episodic retrieval compared to semantic retrieval. Despite a reduced cluster size for PPI-V1$_{EPISEM}$ compared to PPI-LOC$_{EPISEM}$, the clusters fully overlap, both streaking parietal regions and Precuneus. Additionally, for PPI-LOC$_{EPISEM}$, another cluster emerged within the inferior right LOC. Furthermore, increased bilateral functional connectivity with clusters located over the postcentral and supramarginal gyrus (SMG) was measured exclusively during episodic trials and with LOC as seed region. For detailed information see Table S2.

The opposite contrast revealed considerable overlap between semantic and episodic trials for both seed ROIs. A cluster over the right superior LOC, including parts of the angular gyrus, was more strongly connected to both seed regions during semantic than during to episodic trials. Another area that showed substantial voxel overlap for both seed ROIs during semantic retrieval was the bilateral fusiform area. In the right hemisphere, both fusiform clusters extended to the lingual gyrus whereas in the left hemisphere, only the fusiform cluster of the PPI-LOC$_{SEMEPI}$ contrast reached to the angular gyrus. For full report see Table S2.

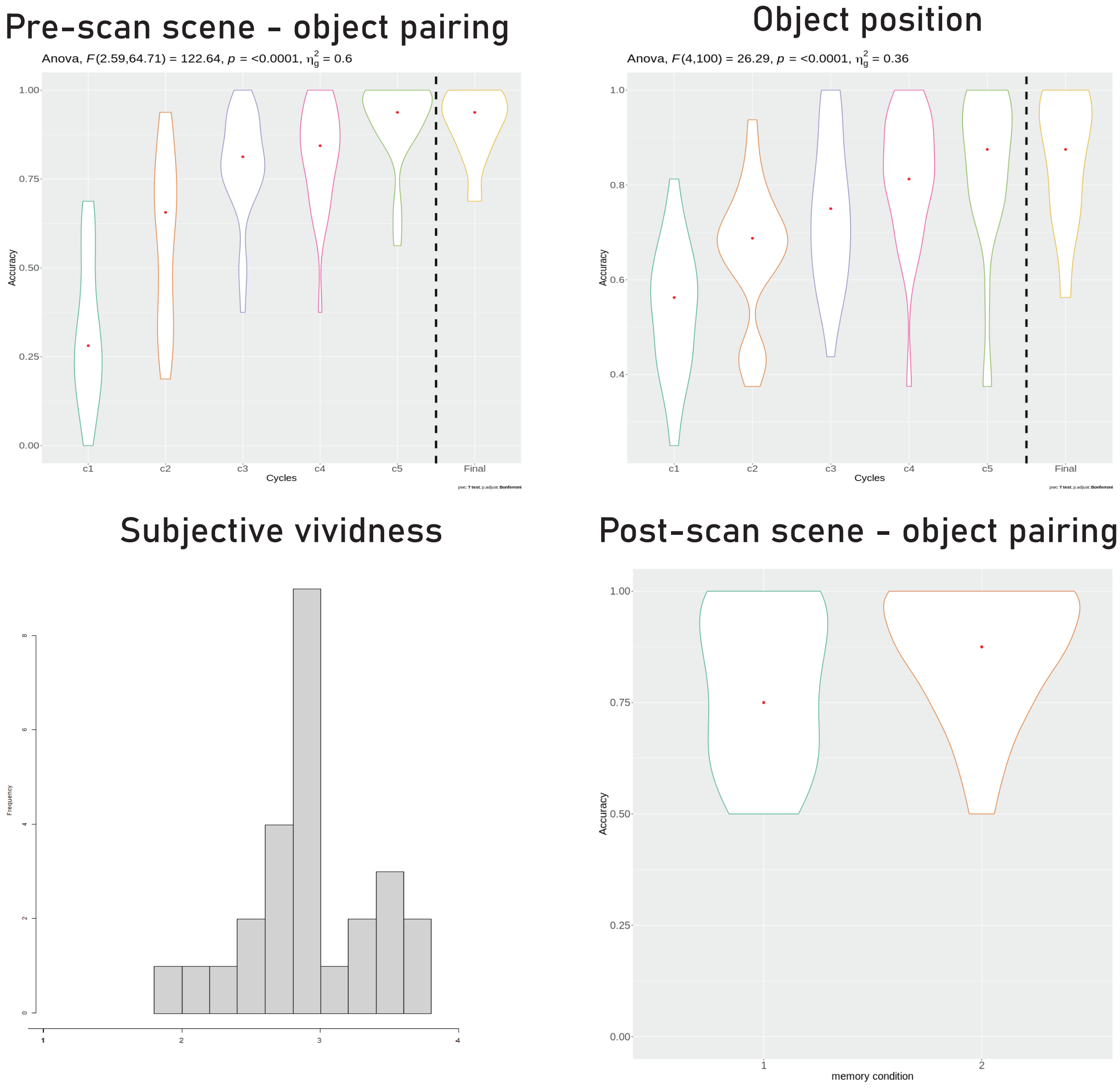


**Fig. S1. Behavioural performance.** (**A and B**) Accuracy for scene-object pairing and for object position over the six learning cycles. (**C**) Histogram of average subjective vividness rating for the retrieved objects during the scanner training phase. Participants used a 4-point Likert scale (1 = “not remembered”, 2 = “remembered but not visualized”, 3 = “visualized with a few details”, 4 = “visualized with a lot of detail”). Participants were encouraged to try to get as many 3s and 4s as possible. (**D**) Post-scan memory test performance for episodic and semantic conditions.

**Table S1. Summary of classification results**

| Classification analysis | Visual field region | ROI | Retrieval condition | Accuracy | Z |
|---|---|---|---|---|---|
| Concurrent | Periphery | V1 | Episodic | .67 | 4.69* |
| | | | Semantic | .64 | 4.58* |
| | | V2 | Episodic | .62 | 4.69* |
| | | | Semantic | .60 | 4.62* |
| | Fovea | V1 | Episodic | .61 | 4.67* |
| | | | Semantic | .60 | 4.61* |
| | | V2 | Episodic | .57 | 4.39* |
| | | | Semantic | .56 | 4.07* |
| Mnemonic | Periphery | V1 | Episodic | .68 | 4.69* |
| | | | Semantic | .67 | 4.67* |
| | | V2 | Episodic | .64 | 4.69* |
| | | | Semantic | .62 | 4.69* |
| | Fovea | V1 | Episodic | .62 | 4.69* |
| | | | Semantic | .61 | 4.58* |
| | | V2 | Episodic | .59 | 4.69* |
| | | | Semantic | .57 | 4.69* |
| Cross-classification | Periphery | V1 | Episodic | .49 | -0.45 |
| | | | Semantic | .50 | 0.43 |
| | | V2 | Episodic | .48 | 0.74 |
| | | | Semantic | .49 | -0.21 |
| | Fovea | V1 | Episodic | .48 | -0.72 |
| | | | Semantic | .49 | -0.26 |
| | | V2 | Episodic | .50 | 0.12 |
| | | | Semantic | .49 | -0.27 |

**Table S1. Summary of classification results.** Sample-averaged accuracies for each classifier type. Asterisks denote significance for the corresponding one-sided signed rank test, $p<.05$.

| ROI | Contrast | Hemisphere | Region (Harvard-Oxford Cortical Structural Atlas) | Z - Local maxima MNI coordinates X | Y | Z | Cluster size (no. of voxels) | Peak *Z* | Cluster-level (FWE-corrected) |
|---|---|---|---|---|---|---|---|---|---|
| V1 | EPI>SEM | | **Cluster 1** | | | | | | |
| | | right | LOC, superior division | 11.5 | -66.5 | 69.5 | 197 | 4.25 | 0.00028 |
| LOC | EPI>SEM | | **Cluster 4** | | | | | | |
| | | right | LOC, superior division | 11.5 | -66.5 | 71.5 | 439 | 4.52 | 8.27e-09 |
| | | | **Cluster 3** | | | | | | |
| | | right | Postcentral gyrus | 31.5 | -32.5 | 43.5 | 139 | 4.33 | 0.00253 |
| | | | **Cluster 2** | | | | | | |
| | | left | SMG; Postcentral gyrus | -50.5 | -30.5 | 41.5 | 134 | 4.05 | 0.0033 |
| | | | **Cluster 1** | | | | | | |
| | | right | LOC, inferior division; middle temporal Gyrus | 53.5 | -62.5 | -0.5 | 119 | 3.93 | 0.00742 |
| V1 | SEM>EPI | | **Cluster 4** | | | | | | |
| | | right | LOC, superior division; Angular Gyrus | 41.5 | -62.5 | 31.5 | 222 | 4.82 | 9.45e-05 |
| | | | **Cluster 3** | | | | | | |
| | | right | Fusiform Cortex; Fusiform Gyrus; Lingual Gyrus | 23.5 | -58.5 | -8.5 | 185 | 3.99 | 0.000478 |
| | | | **Cluster 2** | | | | | | |
| | | left | LOC, inferior division | -44.5 | -82.5 | -8.5 | 110 | 4.45 | 0.0187 |
| | | | **Cluster 1** | | | | | | |
| | | left | Fusiform Cortex | -28.5 | -44.5 | -16.5 | 102 | 4.12 | 0.0287 |
| LOC | SEM>EPI | | **Cluster 3** | | | | | | |
| | | right | Fusiform Cortex; Fusiform Gyrus; Lingual Gyrus | 27.5 | -62.5 | -14.5 | 231 | 4.55 | 3.14e-05 |
| | | | **Cluster 2** | | | | | | |
| | | right | LOC, superior division; Angular Gyrus | 41.5 | -60.5 | 31.5 | 109 | 3.99 | 0.0132 |
| | | | **Cluster 1** | | | | | | |
| | | left | Fusiform Cortex; Fusiform Gyrus; Angular Gyrus | -30.5 | -56.5 | -6.5 | 91 | 4.45 | 0.0374 |

**Table S2. Summary of clusters surviving correction in PPI analysis.**
